# High-throughput, label-free detection of DNA origami in single-cell suspensions using origamiFISH-Flow

**DOI:** 10.1101/2023.10.31.564845

**Authors:** Shana Alexander, Wendy Xueyi Wang, Chung-Yi Tseng, Travis R. Douglas, Leo Y.T. Chou

**Author notes:** Correspondence to: Leo Y. T. Chou Institute of Biomedical Engineering, University of Toronto, Toronto, ON, Canada.

## Abstract

Structural DNA nanotechnology enables custom fabrication of nanoscale devices and promises diverse biological applications. However, the effects of design on DNA nanostructure (DN)-cell interactions in vitro and in vivo are not yet well-characterized. origamiFISH is a recently developed technique for imaging DNs in cells and tissues. Compared to the use of fluorescent tags, origamiFISH offers label-free and structure-agnostic detection of DNs with significantly improved sensitivity. Here, we extend the origamiFISH technique to quantifying DNs in single-cell suspensions, including nonadherent cells such as subsets of immune cells, via readout by flow cytometry. This method, referred to as origamiFISH-Flow, is high-throughput (e.g., 10,000 cells per second) and compatible with immunostaining for concurrent cell-type and -state characterization. We demonstrate that origamiFISH-Flow enhances signal-to-noise ratio by up to 20-fold compared to dye labeling approaches, leading to the capture of >25-fold more DN^+^ cells at low, single-picomolar DN uptake concentrations. We additionally show the use of origamiFISH-Flow to profile cell-type and shape-specific DN uptake patterns across cell lines and splenocytes and quantify in vivo DN accumulation in lymphoid organs. Together, origamiFISH-Flow offers a new tool to interrogate DN interactions with cells and tissues, while providing insights for tailoring their designs in bio-applications.

## Introduction

Nanostructures assembled from DNA are now being investigated for biological applications such as drug carriers^1–3^, vaccines^4,5^, biosensors^6–8^, and cell-instructive biomaterials^9–11^. Many of these applications require controlling the cell uptake or subcellular localization of DNs, such as the delivery of drugs into specific cell types, clustering of cell-surface receptors^12^, or entry into the cytoplasm or nucleus^13^. These applications have motivated studies to understand the effects of DN design such as size, shape, stability, and surface chemistry on their interactions with biomolecules, cells, and tissues^14–18^. However, existing methods of detecting DNs in cells and tissues have limitations. The field standard involves labeling, or tagging, the DN with fluorescent dyes for detection by fluorescence imaging or flow cytometry, although several studies have shown that signals generated from such approaches do not always reliably reveal the localization of DNs, but may instead arise from dissociated dyes^19–21^. Additionally, the labeling capacity of each DN is typically on the orders of tens of dyes^21–23^, thus limiting detection sensitivity, especially for in vivo applications. While it is possible to increase the degree of labeling via further engineering, dye labels may have nonspecific interactions with biological tissues that alter the biological behaviour of DNs^19,24–26^.

origamiFISH is a recently developed imaging technique for visualizing DNs in cells and tissues^27^. Compared to dye labeling, this approach does not require labeled DNs, offers greater detection sensitivity, and can image diverse DN shapes without structure-specific optimization. Briefly, the technique is based on the in-situ hybridization of oligonucleotide “initiator” probes to the scaffold strand of DNs in fixed cells. Each bound initiator serves as a nucleation site for the in-situ growth of two metastable, dye-labeled DNA hairpins into a tethered fluorescent polymer via hybridization chain reaction (HCR, Fig. 1a). Studies in vitro have shown that each initiator probe can generate a >10-kb long polymer, equivalent to >100 hairpins^28^. Given ∼20 probes per DN scaffold, this offers a high degree of signal amplification and sensitivity^27^. In our study, we used “split initiators” formed from a pair of probes binding adjacent sites on the target, which reduces nonspecific binding and improves signal-to-background^27,28^. With confocal microscopy readout, origamiFISH delivers high-content information regarding the spatial distribution of DNs in cells and tissues. However, an inherent limitation of imaging-based methods is the relatively low throughput in data acquisition and quantification. Another limitation is that it is not suited for the analysis of nonadherent cells such as hematopoietic cells or single-cell suspensions isolated from primary tissues.

**Figure 1.**
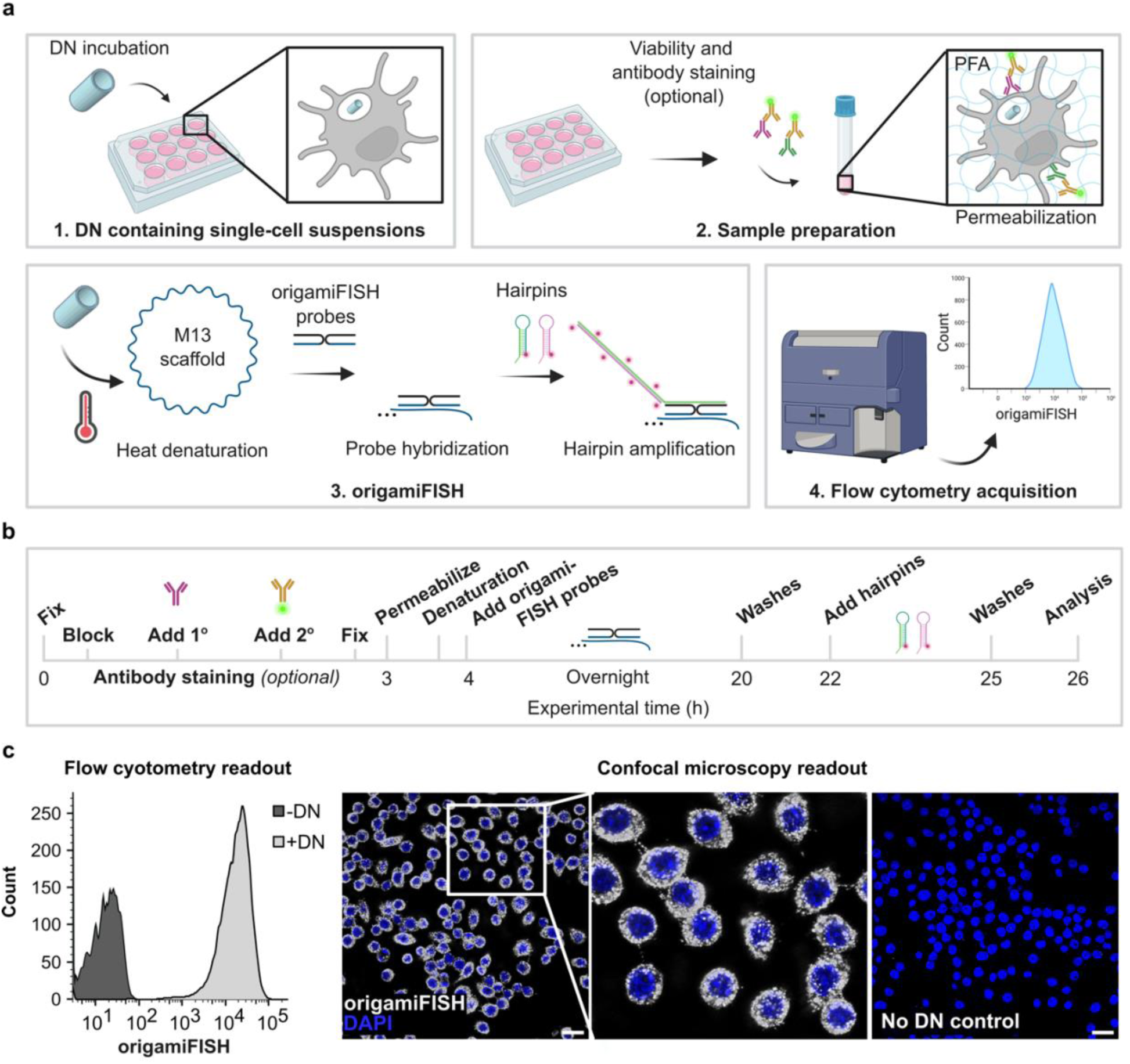
Development of origamiFISH-Flow for high throughput detection of DNs in single-cell suspensions. (a) Overview of the origamiFISH-Flow workflow includes four steps: 1) generate DN-containing, single-cell suspensions, 2) sample preparation, 3) origamiFISH including heat denaturation at 50°C, probe hybridization, and hairpin amplification, and 4) acquisition and analysis on a flow cytometer. (b) Experimental time for origamiFISH-Flow with optional antibody staining. (c) Comparison of origamiFISH signal readout via flow cytometry versus confocal microscopy. In both experiments, RAW264.7 macrophages were exposed to 1 nM DN rectangles for 30 min. Scale bar = 20 *µ*m

Here, we describe an extension of the origamiFISH technique for readout by flow cytometry, referred to as origamiFISH-Flow. This method combines the sensitivity and label-free nature of origamiFISH with the high throughput of flow cytometry. This enables rapid quantification of DNs in single-cell suspensions, either following in vitro cell uptake or tissue dissociation. We demonstrate origamiFISH-Flow has higher sensitivity over traditional dye labeling approaches and use it to profile DN uptake patterns across four cell types. We demonstrate compatibility between origamiFISH-Flow and immuno-staining for simultaneous DN detection and immunophenotyping in splenocytes. Finally, we use origamiFISH-Flow to quantify the trafficking of DNs to the lymph node following subcutaneous injection. Together, we demonstrate a novel tool for analysis of DN interactions with cells and tissues. We envision this technique will enable (1) quantification of DN uptake and biodistribution dynamics in vivo, (2) analysis of the effects of design on DN-cell interactions, and (3) discovery of DN design parameters for cell-and tissue-specific delivery.

## Results and Discussion

### Development of origamiFISH-Flow

The origamiFISH-Flow workflow involves four major steps: (1) generation of single-cell suspensions following cell uptake or tissue harvest, (2) sample preparation including fixation, antibody staining, and permeabilization, (3) in situ labeling of scaffold DNA via origamiFISH^27^, and (4) flow cytometry for signal detection and readout (Fig. 1a). The protocol takes ∼1 day to complete, including a 16-hour overnight probe incubation step and a 3- to 5-hour hairpin amplification step, which can be adjusted based on the degree of signal amplification desired (Fig. 1b). For analysis of primary cells, we also incorporate a cell viability staining step immediately following cell uptake or tissue harvest to filter out any dead cells from the analysis^29^.

Notably, the origamiFISH-Flow workflow differs from the original imaging-based technique in two ways: (1) imaging requires the sample to be on glass coverslips. For in vitro experiments, this means seeding cells on poly-D-lysine (PDL)-treated glass coverslips and mounting the coverslips on microscope slides prior to imaging. In contrast, origamiFISH-Flow is compatible with seeding cells on regular tissue-culture plates and detaching them into suspensions prior to origamiFISH staining; (2) In the imaging-based method, since the cells are adhered to the coverslip, buffer exchanges are performed by pipetting, whereas in origamiFISH-Flow washes are performed via repeated centrifugation and resuspension in buffer.

To establish origamiFISH-Flow, we first asked how the origamiFISH signal from similar samples appear under flow cytometry compared to confocal microscopy. To test this, we incubated RAW264.7 murine macrophages with 1 nM of DN rectangles for 30 minutes and prepared the sample for either origamiFISH imaging or origamiFISH-Flow, following the respective protocols. We selected this condition because it had been tested in the original origamiFISH study, allowing us to benchmark and compare the results. Following uptake, we observed origamiFISH signal appearing as dense, bright puncta under confocal imaging (Fig. 1c), consistent with earlier observations that macrophages efficiently take up DN rectangle at this concentration and timepoint^27^. When assayed by flow cytometry, the sample had a mean fluorescence intensity (MFI) of ∼21,000, or ∼3800× higher than the control sample without DNs (Fig. 1c). These results suggest that our modifications to the protocol allows detection of DNs in cells by flow cytometry.

We next characterized the detection sensitivity of the origamiFISH-Flow method and aimed to adjust parameters to optimize its detection limit. In origamiFISH, the amount of signal is determined by the number of fluorescent polymers bound to each scaffold DNA. The efficiency of this process is tunable by the concentrations of the initiator probes and fluorescent hairpins, as well as by the amplification time. To characterize the effects of these parameters on detection sensitivity, we followed the same cell uptake condition as before, and performed origamiFISH-Flow with probe concentrations varying from 5 to 20 nM. Here, we observed that the MFI remained constant across all probe concentrations tested (Fig. S1), indicating probe quantities can be reduced with minimal loss in signal. In future experiments, we fixed the probe concentration at 10 nM per sample. Next, we tested the effect of amplification time on the signal. Following cell uptake, samples were incubated with fluorescent hairpins for either 1, 3, or 5 hours, with readout via origamiFISH-Flow. Here, we observed a linear relationship between amplification time and signal intensity, which was detectable across all timepoints tested (Fig. S1). Thus, we propose that amplification time can be adjusted to optimize the detection sensitivity for a particular experiment. Lastly, we assayed the effect of hairpin concentration on signal intensity by amplifying samples with either 10, 30, or 60 nM of the fluorescent DNA hairpins. We observed lower signal with 10 and 30 nM compared to 60 nM under 1-hour amplification (Fig. S1). We hypothesize that higher hairpin concentrations can lead to faster nucleation and growth, and that lower hairpin concentrations may require longer amplification time to achieve similar signals, although this remains to be tested. With these parameters, we next assayed the detection limit of origamiFISH-Flow by incubating RAW264.7 cells with DN rectangles across a range of concentrations from 1 pM to 10 nM (Fig. S2). We successfully detected DNs in cells across all concentrations when using 60 nM hairpins and a 3-hour amplification time. We observed an increase in signal with DN concentration, first linearly from 1 to 100 pM, and then rising logarithmically at higher concentrations, suggesting partial signal saturation above the latter concentration range. From these observations, we conclude that origamiFISH-Flow is a sensitive technique for DN quantification, and its dynamic range can be tuned by adjusting the hairpin concentration as well as amplification time.

To enable flexibility in longitudinal experiments, such as studies involving multi-day timepoints, we also evaluated the stability of the origamiFISH signal over time. To this end, we prepared origamiFISH-stained cells and measured their signals after 0, 7, and 14 days of storage at 4°C. We found no statistically significant change in the origamiFISH signal compared to their respective controls (Fig. S2), suggesting the stained samples can be stored for at least two weeks with minimal signal deterioration. To summarize, origamiFISH-Flow offers high sensitivity and tunable dynamic range for DN detection in cells, with excellent signal stability over time.

### Comparison of origamiFISH-Flow and dye-labeling for DN uptake quantification

We next compared the sensitivity of origamiFISH-Flow to existing dye-labeling approaches for detection of DNs in cells. To do this, we functionalized the DN rectangle with 12 copies of Cy3 using the standard “handle–anti-handle” attachment chemistry^30^ and added them to RAW264.7 macrophages for 30 minutes at concentrations ranging from 1 pM to 10 nM. We then assessed cell uptake, either directly via flow cytometry on the Cy3 channel, or first processing the samples through origamiFISH followed by flow cytometry.

We observed interesting differences in DN uptake distributions measured using the two methods. Specifically, origamiFISH showed tighter DN uptake distributions, whereas dye labeling resulted in broader DN uptake distributions across all concentrations, with some concentrations even showing multiple populations (Fig. 2b). The broad distribution in the Cy3 channel raises the possibility that cells are taking up a heterogeneous population of materials, such as dissociated dyes from the structure, although this remains to be tested. Additionally, origamiFISH exhibited higher signal-to-background (i.e. fold-increase in MFI) compared to Cy3 across all concentrations tested, (Fig. 2c). At 1 pM, origamiFISH signal-to-background ratio was ∼2.6-fold higher compared to Cy3. At 10 nM, this difference increased to 19.5-fold. These findings are consistent with our previous findings from confocal imaging, where origamiFISH detected more DN localizations per cell than observed with Cy3 labeling across the same concentration range^27^. Notably, image analysis of the confocal microscopy data was performed on ∼150 cells per condition, compared to ∼25,000 cells in this flow cytometry experiment, suggesting that origamiFISH is also compatible with this detection format.

**Figure 2.**
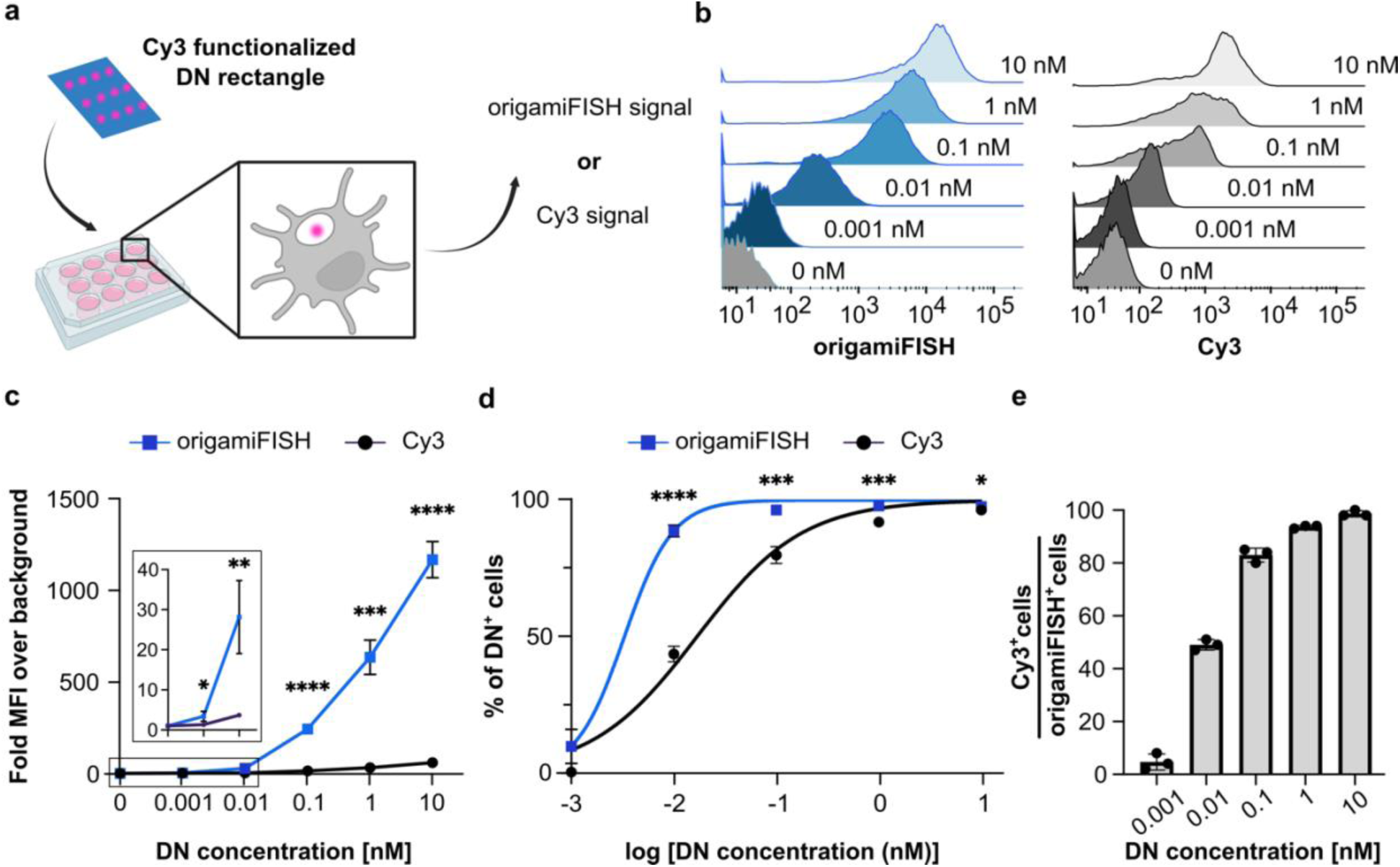
origamiFISH-Flow is more sensitive than dye labeling for DN detection. (a) RAW264.7 macrophages were incubated with DN rectangles labeled with 12 copies of Cy3 for 30 minutes. Uptake was quantified via flow cytometry of the Cy3 channel or the origamiFISH channel following processing. (b) Histograms of the origamiFISH and Cy3 channels at varying DN uptake concentrations. (c) Quantified MFI of origamiFISH and Cy3 signals by normalizing to the respective background signals of each channel. (d) Percentage of DN^+^ cells captured via origamiFISH-Flow or Cy3 fluorescence. The concentration was plotted on log scale and the data were fit to a dose-response curve by least squares (R^2^ for origamiFISH = 0.991 and Cy3 = 0.979). (e) Amount of Cy3^+^ cells as a percentage of origamiFISH^+^ cells across DN concentrations. Error bars represent standard deviation (SD). Statistical significance analyzed with unpaired parametric T test. ****p<0.0001, ***p<0.001, **p<0.01, *p<0.05, ns p>0.05.

We next compared the number of DN^+^ cells captured using either origamiFISH or Cy3 labeling, using the negative control, i.e. 0 nM sample, to establish our gating strategy (Fig. S3). Here we found that origamiFISH detected a higher percentage of DN^+^ cells across all concentrations compared to Cy3 (Fig. 2d). For example, Cy3 labeling detected ∼4%, 50%, and 80% of cells labeled as origamiFISH^+^ at the 1, 10, and 100 pM conditions, respectively (Fig. 2e). Together, this demonstrates the enhanced sensitivity of origamiFISH-Flow compared to dye labeling approaches for detecting DNs in cells.

### Analysis of DN shape-specific uptake in different cell types

After establishing the sensitivity of origamiFISH-Flow, we next applied it to probe the effects of DN shape on cell uptake (Fig. 3). As with other classes of nanomaterials, the shape of DNs is known to modulate their interactions with cells, such as uptake and activation^14,31,32^. Thus, we asked whether origamiFISH-Flow could reveal similar shape-specific uptake patterns as to what has been reported in the literature. We tested three different DN shapes: 1-dimensional (1D) rods, 2D rectangles, and 3D barrels (Fig. 3a, S4). These DNs were incubated with four different cell types: murine macrophages (RAW264.7), bone-marrow derived dendritic cells (BMDCs, Fig. S5), as well as human-derived embryonic kidney 293 cells (HEK293), and Jurkat T cells. In addition to spanning two species of origin, these cells encompass both phagocytes (RAW264.7, BMDC) and non-phagocytes (HEK293, Jurkat), and both adherent (RAW264.7, HEK293) and loosely or non-adherent cells (BMDCs, Jurkat). We reasoned that such diversity of cell types could allow us to validate origamiFISH-Flow for robustness and ensure broad utility. As before, DNs were introduced to cells at a concentration of 1 nM for 30 minutes. Among the cell types evaluated, we observed the highest uptake for RAW264.7 macrophages (Fig. 3b). Across shapes, uptake was the highest for DN rectangle followed by barrel and rod, consistent with results previously obtained by origamiFISH confocal imaging of RAW264.7 cells^27^. Interestingly, this shape-specific preference is consistent across all four cell types tested, suggesting that previous conclusions indicating preferential uptake of more compact structures over elongated ones is generalizable^14^. For example, the 1D rod, which measures 400 nm long and 6 nm in diameter, exhibited the least uptake across all cell types. This is hypothesized to result from the substantial amount of energy required to deform the cell membrane for its engulfment. Interestingly, despite both RAW264.7 and BMDCs being phagocytic cells, RAW264.7 cells demonstrated a significantly higher capacity to internalize DN rectangles, with a ∼31-fold higher MFI (Fig. 3b).

**Figure 3.**
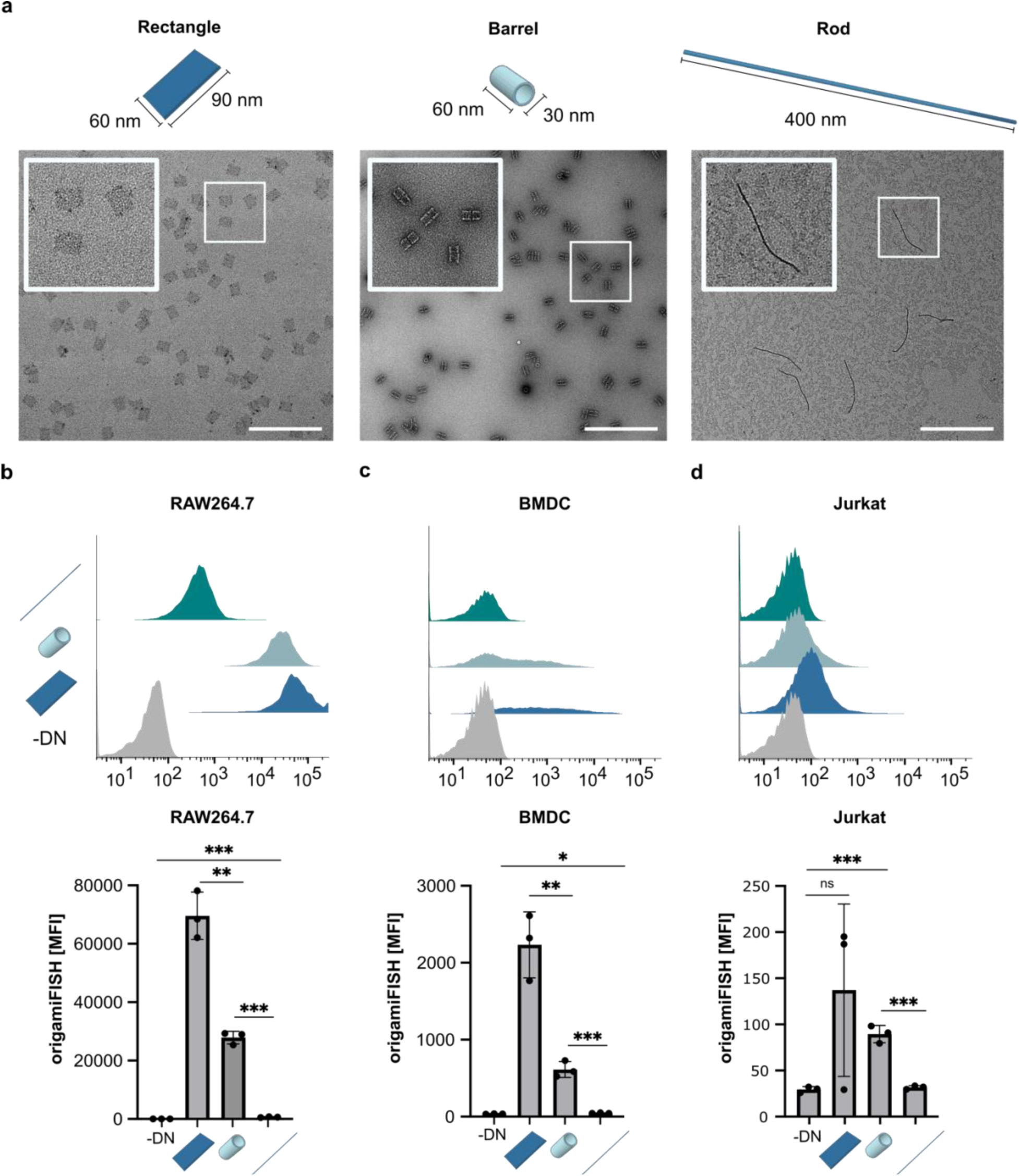
DN shape-specific uptake profiles across cell types assessed by origamiFISH-Flow. (a) Schematic of DN shapes and corresponding transmission electron microscopy images. (b-d) Comparison of origamiFISH-Flow histograms (top) and MFI (bottom) for DN rectangles, barrels, and rods following uptake by (b) RAW264.7 murine macrophages, (c) bone marrow-derived dendritic cells (BMDCs), and (d) human Jurkat T cells. Error bars represent standard deviation. Statistical significance analyzed with unpaired parametric T test. ***p<0.001, **p<0.01, *p<0.05, ns p>0.05. Scale bar = 500 nm

In addition to MFI, we observed differences in DN uptake distribution across cell types. DN uptake by RAW264.7 macrophages exhibited a normal distribution, whereas BMDCs exhibited significantly more widened distribution patterns with a tail towards the higher end of the histogram, suggesting greater heterogeneity in the capacity of BMDCs to internalize DNs compared to RAW264.7 cells. The exact reason for this is unclear but may be due to heterogeneity in the phenotype of primary cells following tissue isolation and ex vivo differentiation. In contrast to the phagocytes, Jurkat cells exhibited minimal uptake of all DN shapes (Fig. 3b), consistent with the notion that T cells are not known to efficiently engulf nanomaterials. Finally, we note that the use of origamiFISH-Flow to assay DN uptake by HEK293 cells was challenging. These adherent cells have a strong propensity to clump, leading to difficulties in cell staining and retaining enough cells following the filtering step required prior to flow cytometry (Fig. S6). Despite these challenges, we observed the highest uptake with DN rectangles, followed by the barrel, and then rod, similar to other cell types. However, we note that more optimization of the cell dissociation protocol may be necessary when dealing with cells or tissues that have the propensity to form tight clumps. Nonetheless, our results demonstrate that origamiFISH-Flow can quantify cell-type and shape-specific uptake profiles of DNs.

### Concurrent immunostaining and origamiFISH-Flow

We next asked whether the origamiFISH-Flow workflow is compatible with immuno-staining, which would enable concurrent analysis of DN uptake with cellular proteins. This capability is useful for assessing DN distribution in mixed cell populations, such as single-cell suspensions harvested from primary tissues.

The origamiFISH-Flow workflow includes an ethanol permeabilization step, which may alter the integrity of cell membrane proteins or the availability of their epitopes for antibody binding. To test this, we first assayed the effect of ethanol permeabilization on dye-conjugated primary antibodies by labeling RAW264.7 cells with either anti-CD11b (PE), anti-CD11c (APC), or anti-F4/80 (FITC), followed by processed these cells through origamiFISH-Flow. We observed that all fluorescent channels exhibited considerable signal loss, with the greatest reduction occurring after ethanol permeabilization (Fig. S7).

This was surprising given we had previously observed excellent signal retention when using Alexa Fluor® (AF)-labeled secondary antibodies to label F4/80 as well as the intracellular proteins EEA1, Lamp1, and pHH3^27^. We hypothesized that the process of permeabilization could alter either the dye or the epitope of the target protein. To first rule out the dye as a factor, we repeated the experiment with an unconjugated primary anti-F4/80 and an AF-labeled secondary antibody. Using this approach, we found the F4/80 signal to be retained (Fig. S7), suggesting the F4/80 epitope remained intact after the origamiFISH protocol, consistent with the original study. However, the FITC dye was unable to persist, consistent with ethanol affecting the fluorescence of the dye.

To investigate whether the use of AF-labeled secondaries with ethanol permeabilization could retain other epitopes, we harvested splenocytes and stained B cells and myeloid cells with anti-CD19 and anti-CD11b antibodies, respectively. Our antibody staining revealed that while the CD11b epitope was unimpacted, there was considerable loss of the CD19^+^ cell population (Fig. S8). This demonstrates that certain epitopes are in fact lost during the protocol when using ethanol as the permeabilization agent. To find a more compatible workflow, we next tested different permeabilization agents, i.e., methanol, acetone^33^ (Fig. S9), Tween 20, and Triton X-100 (Fig. S10), as well as different sequences of workflows (Fig. S10). We found that in general, protein labeling should be performed prior to origamiFISH, and the use of a post-fixation step helped retain the signal, while the optimal permeabilizing reagent that retained antibody signal is protein specific. For example, the use of 0.2%v/v Triton X-100 was effective for retaining signal from antibodies against B220 and CD3, while CD11b was more compatible with ethanol. A summary of the compatibility of permeabilization agents with antibody retention and origamiFISH detection is presented in Table 1. These results underscore the need to validate the workflow for each new antibody panel.

**Table 1.**
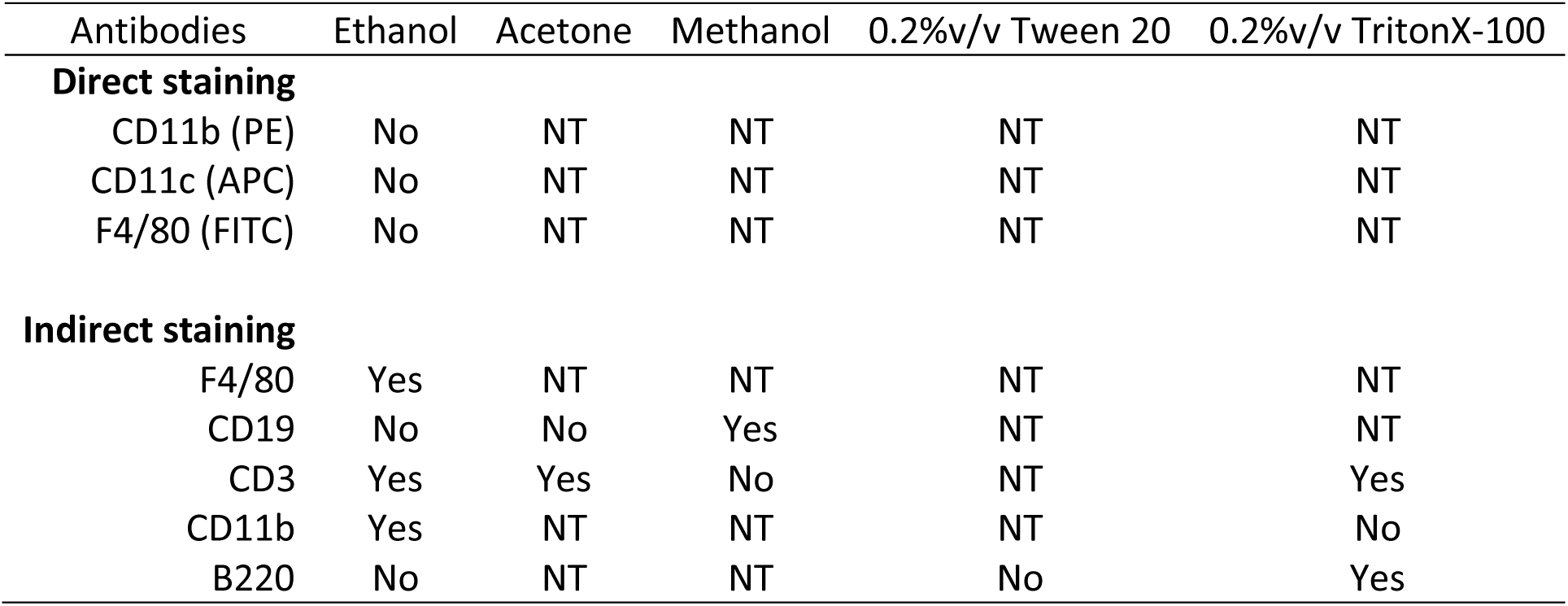
List of antibodies and permeabilization reagents tested for compatibility with antibody staining and origamiFISH-Flow. Indirect immuno-staining was performed using AF594-labeled secondary antibodies against the primary. No, not compatible, Yes, compatible, NT, not tested.

### DN uptake patterns in splenocytes analyzed by origamiFISH-Flow

Using protocols optimized for both antibody staining and origamiFISH, we assayed for DN uptake by splenocytes ex vivo (Fig. 4). Briefly, splenocytes seeded in 12-well plates were introduced to 1 nM of either DN rectangles or buffer for 30 minutes (Fig. 4a). Cells were stained with a viability dye, fixed, processed through antibody staining followed by origamiFISH-Flow, then analyzed immediately. To assess cell-specific uptake patterns, cells were stained with antibodies against B220, a pan-B cell marker, CD3, a T cell marker, and CD11b, a myeloid cell marker. For CD11b staining, we used ethanol as the permeabilization agent, while 0.2%v/v Triton X-100 was used for CD3 and B220 stained cells, which enabled concurrent origamiFISH detection and antibody staining (Fig. 4b). We found that B cells and T cells each accounted for ∼42% of splenocytes, while myeloid cells accounted for ∼4%, consistent with literature (Fig. 4c)^34^. Interestingly, we found that ∼50% and ∼52% of the CD11b^+^ and B220^+^ cells, respectively, were DN^+^ (Fig. 4d). CD11b is a cell-surface integrin protein expressed on many myeloid immune cells including neutrophils, macrophages, and dendritic cells^35^. These cells were expected to internalize nanoparticles^36^, as we observed for RAW264.7 macrophages and BMDCs (Fig. 3). By contrast, B cells are lymphocytes known for secreting antibodies and presenting antigens^37,38^, but are not well-recognized to take up nanomaterials. Across T cells, we found that only ∼9% were DN^+^. This finding was not surprising and corroborated with our observations on Jurkat cells (Fig. 3d). Interestingly, a prior study by Tsoi et al. also demonstrated that both B cells and Kupffer cells (liver-resident macrophages) efficiently interact with quantum dots, another type of nanoparticle^39^. While similar proportions of B cells and myeloid cells were DN^+^, the MFI for CD11b^+^ myeloid cells was ∼2.7× higher than the MFI of B220^+^ B cells (Fig. 4e). This suggests that myeloid cells can internalize more DNs per cell compared to B cells. Tsoi et al. showed a similar finding, with Kupffer cells having MFI ∼1.8 × higher than B cells following exposure to quantum dots^39^. Interestingly, in our study, B cells had the lowest MFI across all three cell types, despite having the largest DN^+^ cell population (Fig. 4e). Together, we showed that the use of origamiFISH-Flow can reveal cell type-specific DN distribution in heterogeneous primary cell populations.

**Figure 4.**
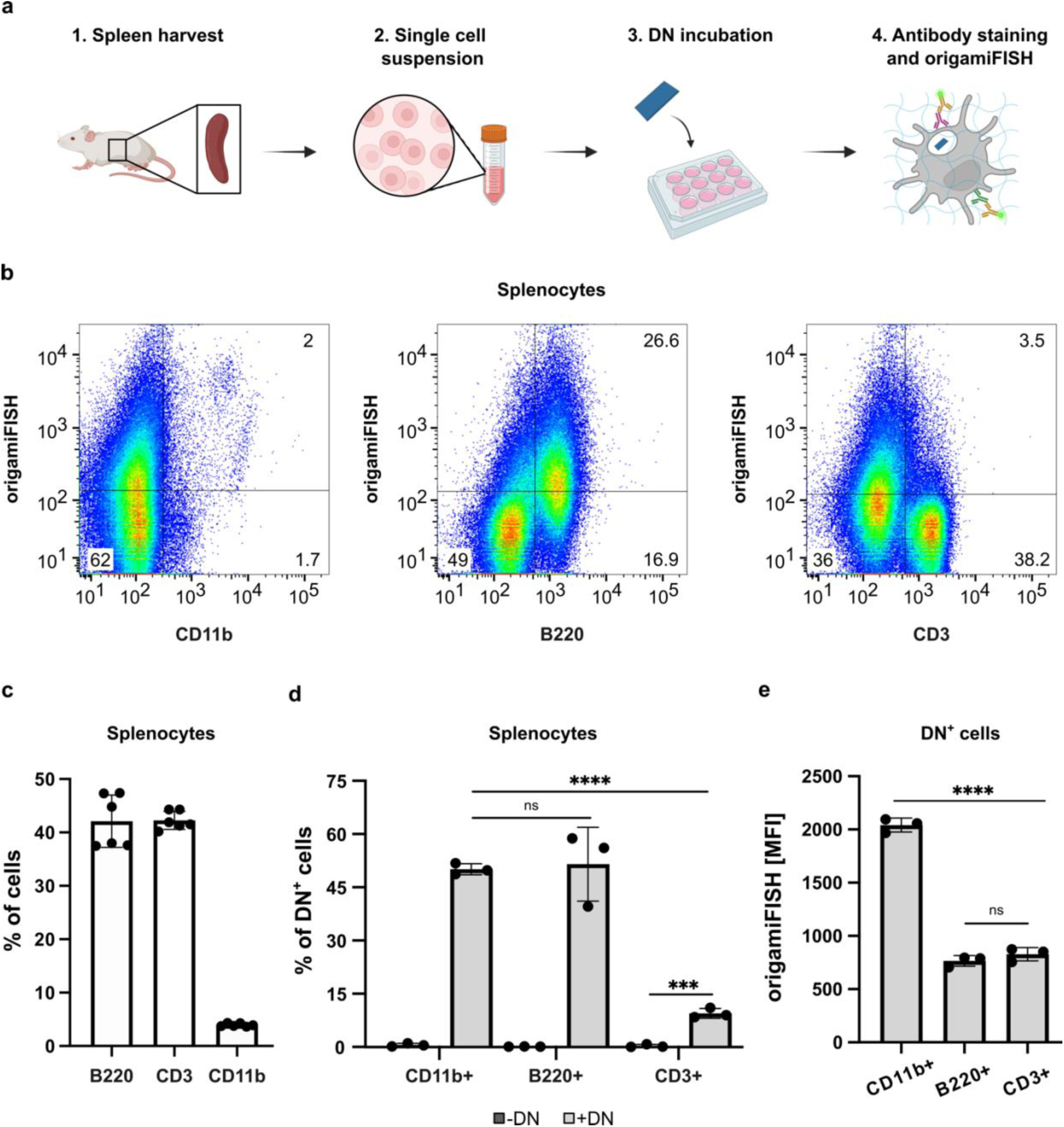
origamiFISH-Flow enables simultaneous detection of DNs and protein markers in mixed cell populations. (a) Overview of experimental workflow. Single-cell suspensions of splenocytes isolated from mice were exposed to 1 nM DN rectangles for 30 minutes. Cells were fixed, separately stained for antibodies against B220, CD3, and CD11b, followed by origamiFISH-Flow. Permeabilization was performed using ethanol for CD11b while 0.2%v/v Triton X-100 was used for B220 and CD3. (b) origamiFISH and antibody staining of CD11b, B220, and CD3 across splenocytes. (c) Proportion of B220^+^, CD3^+^, and CD11b^+^cells in splenocytes. (d) The percentage of DN^+^ cells across CD11b^+^, B220^+^, and CD3^+^ cell populations. (e) MFI of DN^+^ cells across CD11b^+^, B220^+^ and CD3^+^ cells. Statistical significance analyzed with two-tailed unpaired parametric T test. ****p<0.0001, ***p<0.001, ns p>0.05.

### Quantification of in vivo DN accumulation in lymph nodes via origamiFISH-Flow

DN biodistribution is commonly measured with in vivo imaging systems such as the IVIS® using fluorescently tagged nanostructures. However, whole-animal and whole-organ fluorescence imaging provide only semi-quantitative information^40^ due to the poor penetration of light through biological tissues and have limited spatial resolution. Alternatively, biodistribution can be assessed histologically at the expense of bulk-level quantification, or else by dissociating tissue into single-cell suspensions for analysis on flow cytometry. We previously demonstrated using origamiFISH for imaging DNs in tissue sections. We now show that origamiFISH-Flow can quantify DNs in single cells isolated from whole organs for characterizing DN-organ and DN-cell interactions.

To demonstrate the sensitivity and single-cell quantification possible with origamiFISH-Flow for in vivo applications, we evaluated the draining of a DN-based immunomodulatory material to secondary lymphoid organs, which are enriched in immune cells that orchestrate the immune response (Fig. 5a). Nanoparticle size is one parameter that mediates its trafficking within the lymphatic system and systemic circulation. For example, nanoparticles larger than 100nm drain less efficiently than smaller nanoparticles and are instead retained at the injection site or rely on cell-mediated trafficking to reach the lymph nodes^41,42^. While DNs are increasingly being investigated as a platform to build vaccines^4,5,11^, the design parameters for their delivery to lymphoid tissues have not been characterized in detail. Here we assess the ability of DN rectangle, which has dimensions of 60nm×90nm, to accumulate in the draining lymph node and spleen over time (Fig. 5a). These DN rectangles were modified to display twelve copies of CpG oligonucleotides, which are pathogen-associated molecular patterns that have clinical relevance as vaccine adjuvants, spaced ∼14 nm and ∼22 nm apart in a rectangular array (Fig. 5b). CpG motifs are known to activate Toll-like receptor 9 (TLR9) expressed in various immune cells^43^ and promote their secretion of proinflammatory cytokines and chemokines, as well as the upregulation of activation markers^43^. We confirmed successful functionalization by incubating the DNs with fluorescently labeled anti-CpG probes and resolving their co-localization by agarose gel electrophoresis (Fig. S11). Functionalized structures were coated with the polymer oligolysine-poly(ethylene glycol), or K10-PEG, at a nitrogen-to-phosphate ratio of 1:1 for increased in vivo stability, as previously reported^44^. The DNs were injected subcutaneously into the left and right tail base of mice at a dose of 50 nM of 100 uL each. At 2-, 4-, 12-, and 24-hours post-injection, the mice were sacrificed, and the draining inguinal lymph nodes from both injection sites as well as the spleen were harvested (Fig. 5c,d). Inguinal lymph nodes were pooled together during dissociation for analysis to reduce biological variability from any single injection site.

**Figure 5.**
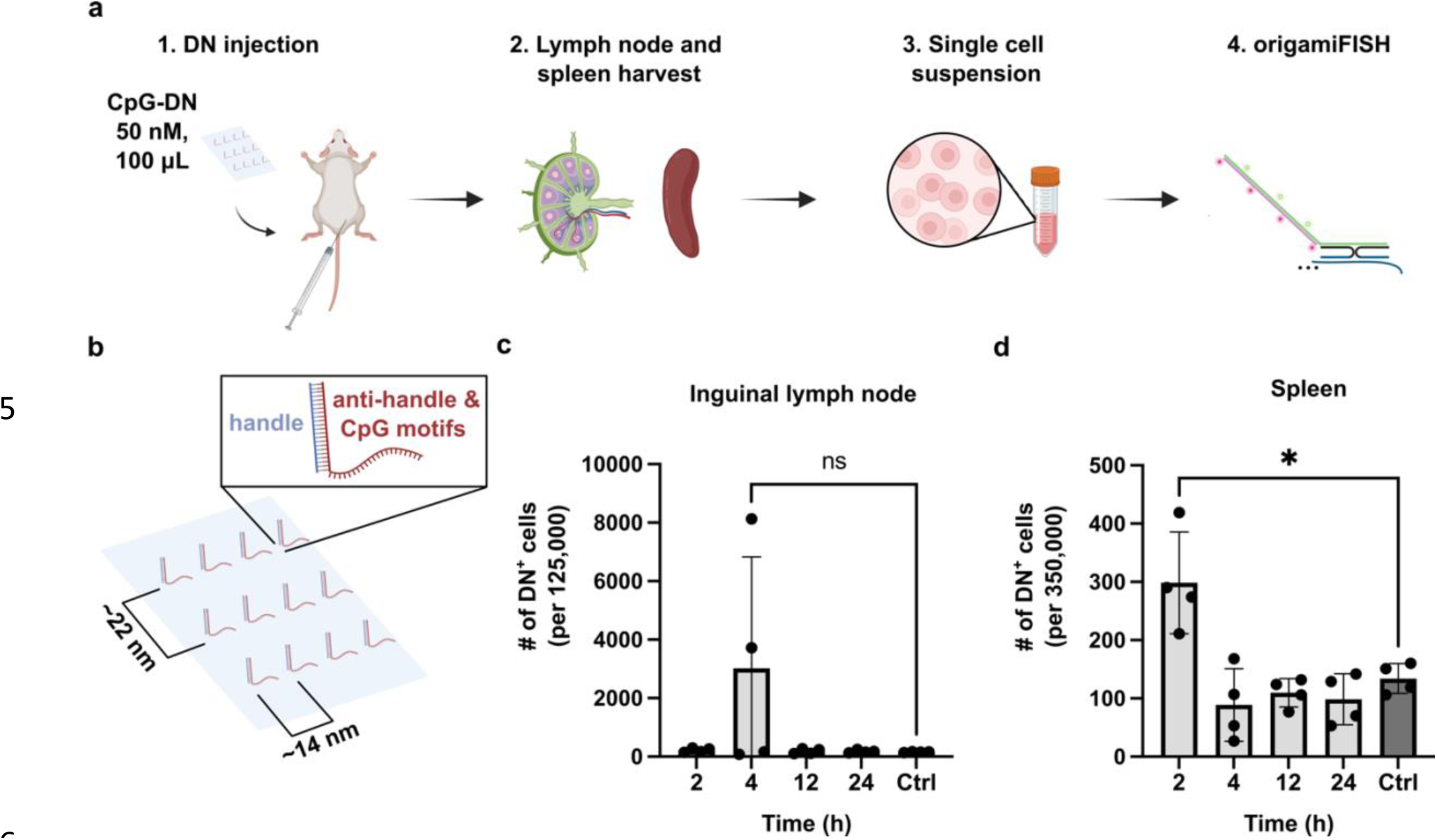
In vivo DN accumulation in lymphoid organs quantified by origamiFISH-Flow. (a) In vivo workflow: 50 nM of 100*µ*L CpG-functionalized DN rectangles (CpG-DN) were injected into the left and right base of the tail. At 2-, 4-, 12-, and 24-hours post-injection, inguinal lymph nodes and spleens were harvested and dissociated, processed by origamiFISH-Flow and analyzed to quantify DN accumulation. (b) Schematic showing the position of the twelve CpG strands on the DN rectangle. (c-d) The number of DN^+^ cells in (c) inguinal lymph nodes and (d) spleen over time compared to naïve control (Ctrl)(n=4). Error bars represent standard deviation. Statistical significance analyzed with unpaired parametric T test. *p<0.05, ns p>0.05.

Our observations with origamiFISH-Flow indicate that the trafficking of DNs to the lymph nodes was variable compared to their systemic distribution to the spleen (Fig. 5c,d). In two of the four mice, we observed a notable increase in lymph node accumulation 4 hours after injection, with DN^+^ cells making up 3.2% to 6.5% out of 125,000 cells analyzed. However, in the other two animals, the origamiFISH signal was comparable to non-injected control mice at the same time point. Across all four replicates in the sample group, the average percentage of DN^+^ cells were ∼2.4%, compared to ∼0.12% in the control group. Despite the low amount of DN^+^ cells, the kinetics of DN drainage is consistent with another study using a 3D square block DN, which also showed peak accumulation in draining lymph nodes at 2 and 4 hours^4^. Liu et al. analyzed the lymph node accumulation of a DN-based vaccine composed of a single-layered DN cylinder. The DN was labeled with Cy5 at the tail-base and injected subcutaneously. They harvested the tissues at 24 hours post-injection and observed signal in the inguinal lymph nodes^5^. In contrast, we detected minimal DN accumulation at this time, using half of the dose compared to Liu et al. Notably, both prior studies performed ex vivo optical imaging of bulk lymph nodes by IVIS®, and therefore did not provide quantitative, single-cell data.

Interestingly, the spleens of the same 4 mice exhibited a consistent number of DN^+^ cells across the time points (Fig. 5d). Specifically, we observed rapid but minimal trafficking to the spleen 2 hours after injection. The average number of DN^+^ cells was ∼2.2-fold higher than the control, consistent with literature demonstrating minimal DN trafficking to the spleen at this time^4,22^.

Together, we demonstrate an application of origamiFISH-Flow for quantifying DN trafficking and biodistribution in vivo. Our data suggests modest draining of CpG functionalized-DN rectangles to the lymph nodes following subcutaneous injection, with minimal systemic distribution to the spleen over a 24-hour window. The use of origamiFISH-Flow enabled the detection of minimal DN accumulation at the single cell-level, previously not possible with whole-animal imaging techniques, and at injection doses difficult to detect via dye labeling. Additionally, our findings motivate the need for more detailed understanding of the biodistribution of DNs, as well as their design principles for delivery to lymph nodes as vaccine platforms.

## Conclusions

This study describes the development and validation of origamiFISH-Flow, a flow cytometry-based approach for label-free detection of DNs in single-cell suspensions. We demonstrated the ability to detect DNs in cells across a range of exposure concentrations from 1 pM to 10 nM, suggesting the method has high sensitivity. We proved that origamiFISH could detect more DN^+^ events compared to traditional dye labeling approaches. Furthermore, we showed that the origamiFISH signal is robust, as the fluorescence intensity within the sample was retained for up to 2 weeks of storage, providing flexibility for multi-day experiments. By applying origamiFISH-Flow to assay effects of DN shape and cell type on DN uptake, we demonstrate the versatility of this method. We further adapted origamiFISH-Flow to be compatible with immunostaining, enabling simultaneous detection of protein expression and DN uptake, although we note the need to validate the sample preparation process for each antibody clone used. This adaptation allowed us to discover the distribution of DNs within splenocytes. Lastly, we used origamiFISH-Flow to quantify the in vivo trafficking of DNs to lymphoid organs. Together, our findings collectively highlight the potential of origamiFISH-Flow for diverse biological applications involving DNs, including in vivo biodistribution studies. The high sensitivity of origamiFISH-Flow will be useful for detecting DN distribution in vivo, especially at lower injection doses and uptake by rare cells, thereby revealing new DN biology as well as guiding the design of DNs for tissue- and cell-type specific delivery.

## Methods

### DN folding

DNs were folded according to previous protocols^14^ with a ThermoFisher ProFlex PCR. ssDNA staples were purchased from IDT. Rectangle DNs were folded with 10 nM of M13mp18 ssDNA scaffold (Bayou Biolabs, P-107) and annealed in 20x excess staples: 80°C for 5 minutes, 60-4°C for 3.16 minutes per °C (for a total of 3 hours). Folding buffer contained 1x modified TE buffer (5 nM Tris, 1 mM EDTA pH 8) with 12.5 mM MgCl_2_. DN barrels were folded with 10 nM of p7308 ssDNA scaffold (Guild Biosciences, D441-020-1mL) and annealed in 10x excess staples: 65°C for 15 minutes, 50-40°C for 98 minutes and 11 seconds per °C (for a total of 18 hours). Folding buffer contained 1x modified TE buffer with 10 mM MgCl_2_. DN rods were folded with 10 nM of p7308 scaffold and annealed in 10x excess staples: 80°C for 5 minutes, 65-25°C (for a total of 18 hours). Folding buffer contained 1x modified TE buffer with 8 mM MgCl_2_.

### DN purification

Rectangle: 10 nM of DN rectangle was combined in a 1:1 v/v with 2X PEG stock consisting of 15%w/w polyethylene glycol 8000 (BioShop, Cat#PEG800), 1X TE buffer, 500 mM NaCl, and 12.5 mM MgCl_2_. The resulting mixture was incubated at room temperature for 10 minutes. Subsequently, the sample underwent precipitation through centrifugation at 14,000g for 30 minutes at 4°C, and the supernatant was carefully removed using a pipette. DN pellet was resuspended in 1/4 of the original crude sample volume, thoroughly mixed, and further incubated in a thermoshaker (Eppendorf ThermoMixer® C) at 30°C and 350 RPM for 20 minutes to facilitate resuspension. The purification was repeated twice. For Cy3 functionalization, Cy3 anti-handles were incubated in 8x excess the number of sites with folded DNs for 1 hour at room temperature after the first purification round. The excess was removed by two additional rounds of purification.

Barrel: 10 nM of DN Barrel was combined in a 1:1 v/v with 2X PEG stock consisting of 14%w/w polyethylene glycol 8000 (Bioshop, Cat#PEG800), 1X TE buffer, 250 mM NaCl, and 10 mM MgCl_2_. The resulting mixture was incubated at room temperature for 30 minutes. Subsequently, the sample underwent precipitation through centrifugation at 16,000g for 40 minutes at 25°C, and the supernatant was carefully removed using a pipette. After re-spinning the sample at 16,000g for 5 minutes at 25°C, any remaining supernatant was also removed with a pipette. The pelleted origami was resuspended in 1/4 of the original crude sample volume, thoroughly mixed, and further incubated in a thermoshaker at 30°C and 500 RPM for 30 minutes to facilitate resuspension. The purification was repeated twice.

Rod: 10 nM of DN rod was combined in a 1:1 v/v with 2X PEG stock consisting of 10% w/w polyethylene glycol 8000 (Bioshop, Cat#PEG800), 1X TE buffer, 250 mM NaCl, and 8 mM MgCl_2_. The resulting mixture was incubated at room temperature for 30 minutes. Subsequently, the sample underwent precipitation through centrifugation at 16,000g for 40 minutes at 25°C, and the supernatant was carefully removed using a pipette. After re-spinning the sample at 16,000g for 5 minutes at 25°C, any remaining supernatant was also removed with a pipette. The pelleted origami was resuspended in 1/4 of the original crude sample volume, thoroughly mixed, and further incubated in a thermoshaker at 30°C and 500 RPM for 30 minutes to facilitate resuspension. The purification was repeated twice.

### DN coating with K10-PEG

All experiments use K10-PEG coated DNs. DNs were incubated for at least 30 minutes at room temperature with methoxy-poly(ethylene glycol)-block-poly(L-lysine hydrochloride (mPEG5K-b-PLKC10, Alamanda polymers, Cat#050-KC010) at a ratio of 1:1 nitrogen in amines:phosphates in DNA.

### Agarose gel electrophoresis

DNs were characterized using 2%w/v agarose gel electrophoresis. Gel was casted by the addition of 2.4 g agarose (FroggaBio, Cat#A87) into 120 mL of 0.5x TBE. The mixture was heated in a microwave and allowed to cool, then mixed with 1.2 mL of MgCl_2_ and 5 uL of SYBR™-Safe (Invitrogen, S33102), and casted into the gel tray. Samples were run on ice at 60V for 2 hours.

### Negative stain transmission electron microscopy

DNs were diluted to 1 nM in 1x folding buffer (based on their shape, previously described) and loaded onto a plasma-cleaned and carbon coated, 400 mesh Ted Pella formvar stabilized with carbon transmission electron microscopy grid (thickness 5-10 nm SFR, Cat#01754-F). The sample was allowed to settle on the grid for 2.5 minutes, followed by the removal of excess liquid using filter paper. Subsequently, the grid was washed with 1x folding buffer, and then stained for 30 seconds with 2% uranyl acetate. The samples were then imaged using a Talos L102C transmission electron microscope.

### Poly-D-lysine (PDL)-treated coverslip preparation

PDL-treated coverslips were prepared according to published protocols^27^. Briefly, glass coverslips (12 mm #1.5 Fisher Scientific cat #NC1129240) were washed with 1 M HCl over night at room temperature, followed by washes with water. They were then submerged overnight at room temperature in 0.1 mg/mL of PDL (Sigma; cat# P0899-50MG). Five more washes were then performed with 18.2 MΩ·cm Milli-Q® water. Coverslips were stored in water at 4°C until use. PDL-treated coverslips were stored for up to four weeks prior to use.

### Probe design

Probes were designed in house following previous protocols^27^.

### Cell culture and uptake

BMDCs were harvested from 6- to 8-week-old C57/B6N mice and either used immediately or cryopreserved. Cells were cultured with 20 ng/mL GM-CSF (Biolegend, 576302) for 7 days with DMEM (5% pen/strep, 10% FBS). A control flow tube of cells was stained with APC anti-CD11c (Biolegend, 117310) to validate differentiation. Cells were seeded 24 hr before the experiment in 12-well plates at densities required to achieve 80-90% confluency (RAW 264.7: 200,000 cells, HEK293: 300,000 cells, Jurkat: 400,000 cells and BMDCs: 500,000 cells). BMDCs were seeded in round bottom 96-well plates, whereas all other cells were seeded in flat-bottom 12-well plates. DNs were incubated with cells in 400 uL of media (DMEM 1X with 1.5 g/L sodium bicarbonate and sodium pyruvate (Wisent Bioproducts, 319-007) for RAW264.7 cells, BMDCs, and HEK293 or RPMI 1640 (ATCC modification) (ThermoFisher, A1049101) (for Jurkat cells complete with 5% pen/strep and 10% FBS) at the reported final concentrations. Uptake occurred for 30-minutes in an incubator at 37°C at 5%CO_2_. Adherent cells were harvested with a cell scraper. Cells were transferred from the well plate into polystyrene flow cytometry tubes. All buffer exchanges are done by centrifuging the tubes at 500g for 5 minutes at 4°C and pouring the supernatant out. Flow tubes are resuspended by flicking.

### Antibody staining and dead cell exclusion

Live and dead cell staining was done prior to antibody staining by adding 1 uL of viability dye (LIVE/DEAD^TM^ Fixable Violet, ThermoFisher, L34964) into 1 mL of the cell solution in 1x phosphate buffer saline (PBS) (Gibco, 14190144) and stained for 30 minutes at room temperature. Following staining, cells were fixed using 4% paraformaldehyde for 10 minutes on the rocker (VWR analog rocker 10127-876). Fixed cells were washes once with PBS. To reduce non-specific binding, antibody blocking was performed with 4% normal donkey serum in 0.1%v/v Tween 20 in PBS for one hour at room temperature. Following blocking, primary antibodies were added to the blocking solution for 20 minutes while rocking (VWR analog rocker 10127-876), in the dark and on ice. The following primary antibodies were used: F4/80 (Biolegend, 123107), CD11b (Biolegend, 117310), CD11c (Biolegend, 117310). Unconjugated antibodies were incubated for 1 hour (1:100) on ice in the dark (Rat IgG2a (Biolegend, 400501), Rat IgG2a (Biolegend, 400601), CD3 (Biolegend, 100202), CD11b (Biolegend, 101202), B220 (Biolegend, 103201). Cells were then stained with Alexa Fluor ® 594-conjugated mouse anti-rat IgG (Jackson ImmunoResearch, 212-585-104) for 30 minutes on ice and in the dark. Antibody-labeled cells are washed twice using FCSB, and post-fixed with 4% paraformaldehyde for 10 minutes on the rocker and washed again with 1X PBS.

### origamiFISH-Flow

Flow tubes containing single-cell suspensions were fixed with 4% PFA for 10 minutes at room temperature on the rocker, followed by a PBS wash. Cells were then permeabilized with ethanol or 0.2%v/v Triton X-100 in PBS for 30 minutes on the rocker, at room temperature. At this step, the cells can be stored in ethanol at −20°C for up to one year. Cells were then washed with 1x PBS. To denature DNs, cells were introduced to Probe Hybridization Buffer (Molecular Instruments) and placed in a water bath at 50°C for 10 minutes. HCR split-initiator probes were then added to the tubes to achieve a final concentration of 10 nM. Probe hybridization occurred for 16 hours at 35°C. Cells were washed three times via incubation in Probe Wash Buffer (Molecular Instruments) for 15 minutes each at 37°C. Cells were then washed twice with 2x saline-sodium citrate (SSC) with 0.1%v/v Tween 20 (0.1% Tween 20 in 2XSSC using diluted 20XSSC in water (ThermoFisher, 15557044)) for 10 minutes at room temperature on a rocker. Fluorescence signal amplification was generated using HCR hairpins. Hairpins were prepared by heating at 95°C for 90 seconds and then cooled to room temperature for 30 minutes in the dark. Hairpins were then resuspended in Amplification Buffer (Molecular Instruments) at a final concentration from 10 to 60 nM and placed into the cell solution. Amplification proceeded from 1 hour to 5 hours, followed by 3x washes with 2x SSC containing 0.1% v/v Tween 20. Following the last wash, cells were resuspended in 500 *µ*L of FCSB and filtered through a 40*µ*m cell strainer to remove any cell aggregates. Samples then proceeded to flow cytometry.

### Mouse strains

All experiments were carried out in accordance with the Canadian Council on Animal Care guidelines for use of animals in research. C57/B6 mice (6-8 weeks) were purchased from Charles River.

### Splenocyte isolation

Spleens were harvested from 6- to 8-week-old C57/B6 mice purchased from Charles River. Mice were sacrificed and spleens were removed and placed in 5 mL of Hanks Balanced Salt Solution (ThermoFisher, 24020117). Using small razors, the spleen was minced and pushed through a 70 *µ*m filter into a 50 mL Falcon tube. Using a plunger from a syringe, the spleen was mashed through the filter and rinsed with 1x PBS. Solution was centrifuged for 5 minutes at 500g at 4°C. Following supernatant removal, samples were incubated with 5 mL of 1x RBC lysis buffer (Cell signaling, 46232S) for 5 minutes on ice to remove red blood cells. 10 to 20 mL of cold 1x PBS was added to the cells and centrifuged again to remove the lysis buffer. RPMI 1640 (ATCC modification) was added to the pellet and cell concentration was quantified with a hemocytometer. For cell uptake, cells were seeded in flat-bottom 12-well plates at a concentration of 5 million cells and then incubated with 1 nM of DNs for 30 minutes at 37°C. 1 million cells were then divided into individual tubes for cell-type specific analysis. origamiFISH was performed with 10 nM of probes, 30 nM of hairpins and 3 hours of amplification, following protocol above. 200,000 live cells were collected per condition.

### Quantification of lymph node and spleen draining

CpG strands with phosphorothioate backbones were purchased from IDT and functionalized to the DN rectangle through 12 handle-anti-handle interactions. DN rectangles were folded in 10x excess staples and then 10 nM of purified DN rectangles were incubated 10x excess CpG overnight at 4°C. Excess CpG was purified with 2 rounds of PEG, as described previously. Fluorescent anti-CpG probes (probes with complementary sequences) were used to validate CpG functionalization with gel electrophoresis. C57BL/6 mice received subcutaneous injection at the left and right tail-base of CpG functionalized rectangle DNs (50 nM of 100 uL) or were naïve (n=4). Mice were sacrificed at their specified time point and the inguinal lymph nodes and spleens were harvested. Tissues were dissociated by mechanical mincing with razors and mashed through a 70 *µ*m filter into a 50 mL falcon tube. Tubes were then spun at 500g for 5 minutes and the supernatant was discarded. Splenocyte pellets were resuspended in 1X RBS lysis buffer as described earlier. Cells then proceeded through origamiFISH-Flow with 5 hours of HCR amplification using AF647 hairpins. Lymph nodes and spleens were analyzed with 125,000 and 350,000 single cells, respectively. origamiFISH was done with 10 nM of probes, 60 nM of hairpins, and 5 hours of amplification.

### Flow cytometry

Flow cytometry was carried out on the LSR Fortessa using a minimum of 10,000 events and data was analyzed using FlowJo.

## Author Contributions

S.A. and L.Y.T.C. designed the experiments. S.A. executed all experiments and analysis with the help of W.X.W. conducting origamiFISH with confocal readouts. TEM images were provided from T.R.D.. C.Y.T. contributed to project direction and synthesis of CpG-functionalized DNs. S.A. and L.Y.T.C. wrote the manuscript with input from all authors.

## Acknowledgments

S.A. acknowledges the Ontario Graduate Scholarship, the SCACE Graduate Fellowship, the STARITA Graduate Fellowship, and the FASE Graduate Student Endowment Fund for support. W.X.W. acknowledges the PRiME initiative for fellowship support. C.Y.T. acknowledges the SCACE Graduate Fellowship, CALLUM Memorial Fellowship, and the STARITA Graduate Fellowship. T.R.D. acknowledges the Precision Medicine Initiative (PRiME) fellowship. L.Y.T.C. acknowledges NSERC Discovery Grant, Medicine by Design, and the Canadian Foundation for Innovation for funding. Some of the figures were made with BioRender.

