## Supplemental Information for "High-throughput, label-free detection of DNA origami in single-cell suspensions using origamiFISH-Flow"

### Supplementary Figures:

**Fig. S1:** origamiFISH-Flow parameters can be altered to optimize DN detection

**Fig. S2:** origamiFISH-Flow detects DNs over a wide range of concentrations and exhibits high signal stability

**Fig. S3:** The gating strategy for RAW246.7 macrophages with uptake of 1 pM to 10 nM of DN rectangles

**Fig. S4:** Characterization of DN rectangle, barrel, and rods assessed through 2% agarose gel electrophoresis

**Fig. S5:** Validation of BMDC differentiation using antibody staining

**Fig. S6:** The gating strategy of the HEK293 cells after uptake of DN shapes

**Fig. S7:** Flow cytometry data of the signal loss using conjugated primary antibodies during origamiFISH-Flow protocol

**Fig. S8:** Signal loss and retention of certain epitopes on splenocytes after the origamiFISH-Flow protocol

**Fig. S9:** Effects of replacing fixation and ethanol permeabilization steps with methanol or acetone as an alternative to ethanol for origamiFISH detection

**Fig. S10:** Workflow optimization to combine antibody and origamiFISH-Flow staining protocols

**Fig. S11:** 2% Agarose gel electrophoresis of CpG functionalized rectangle DN used for in vivo experiment

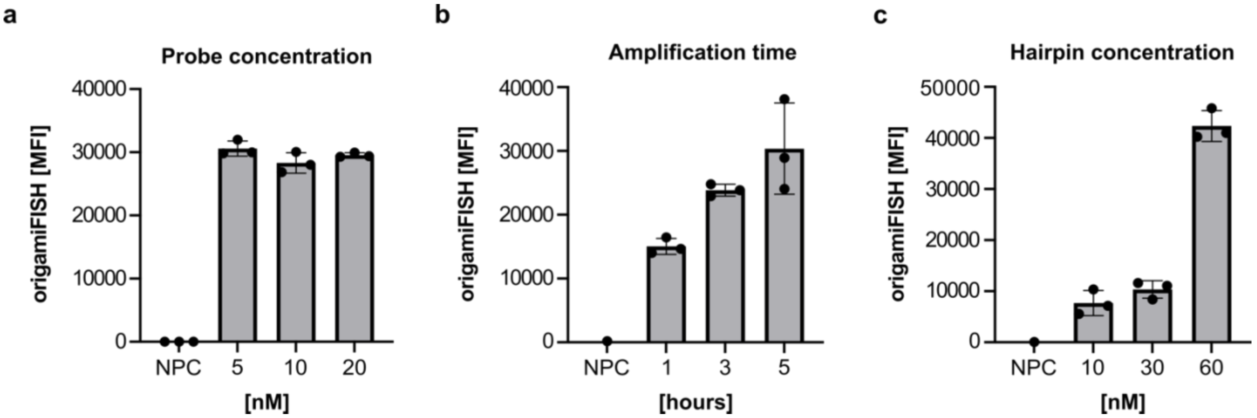


**S1.** origamiFISH-Flow parameters can be altered to optimize DN detection. (a) Uptake of DN rectangles in RAW264.7 cells for 30 minutes. origamiFISH signal remains constant using 5, 10, or 20 nM of probes and 10 nM of AF647 hairpins with 1-hour amplification. (b) origamiFISH signal can be increased by altering the amplification time from 1 hour to 5 hours, using 10 nM of probes and 60 nM of AF647 hairpins. (c) With 10 nM of probes and 1 hour amplification time, origamiFISH signal can be detected with 10, 30, or 60 nM of AF647 hairpins.

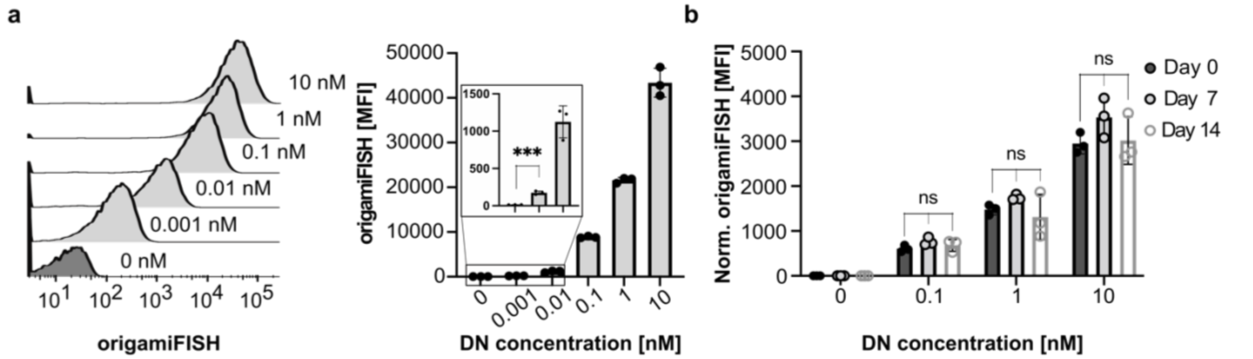


**S2.** origamiFISH-Flow detects DNs over a wide range of concentrations and exhibits high signal stability. (a) Histograms (left) and corresponding bar plots (right) of origamiFISH-Flow signal following 30-minute uptake of DN rectangles from 1 pM to 10 nM in RAW264.7 macrophages. (b) origamiFISH-Flow signal measured on day 0, 7, and 14 from samples in panel (a). Normalized origamiFISH signals were calculated by dividing the MFIs of each sample by the control to account for day-to-day instrument variability. Statistical significance was performed using a two-tailed, unpaired T test for (a) and a one-way ANOVA in (b). *** p<0.001, ** p<0.01, * p<0.05, ns p>0.05.

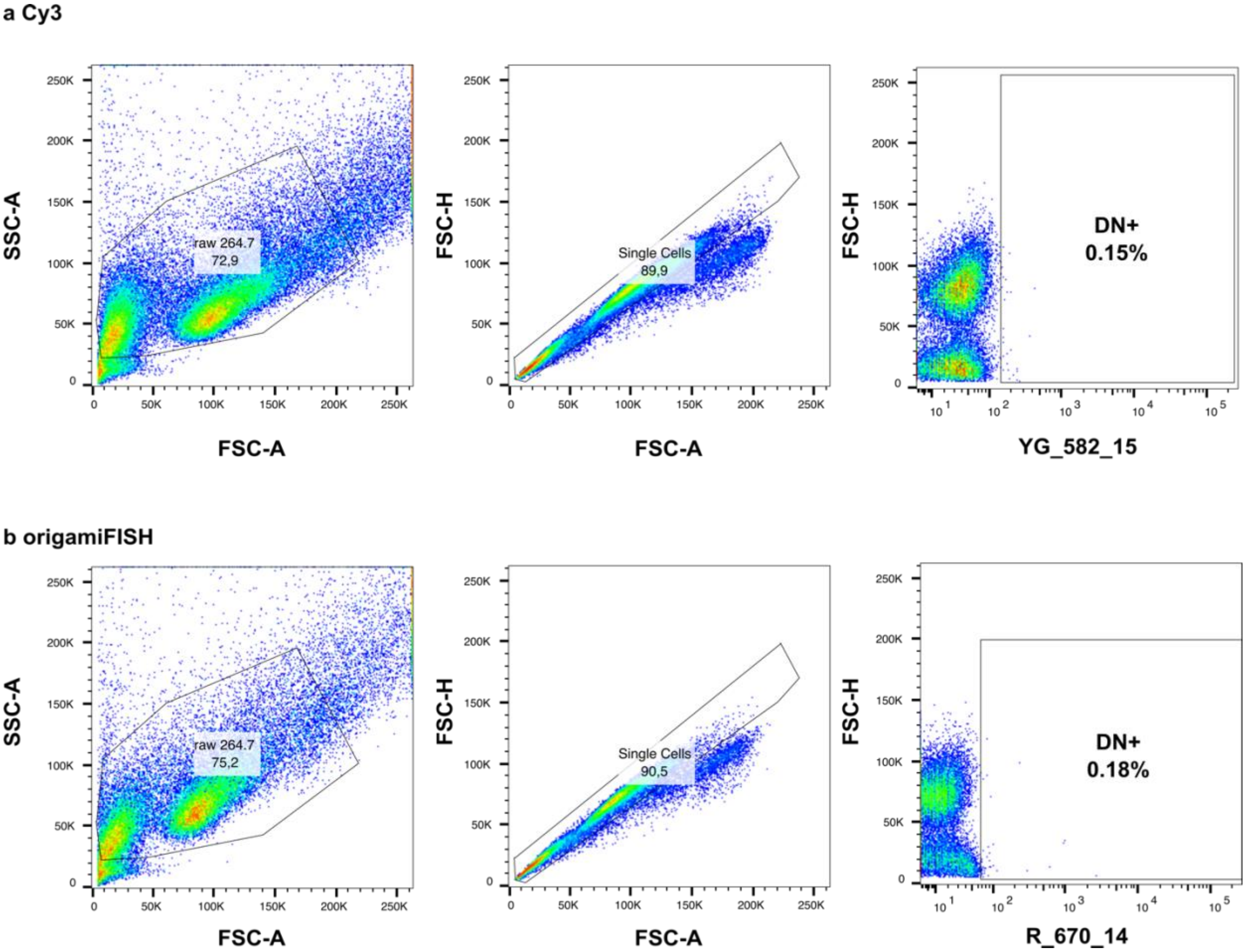


**S3.** The gating strategy for RAW246.7 macrophages with uptake of 1 pM to 10 nM of DN rectangles. The 0 nM control sample was gated on single cells to draw the DN$^+$ gate. This gate was applied to the other conditions in both the (a) the Cy3 channel and (b) the origamiFISH channel (AF647).

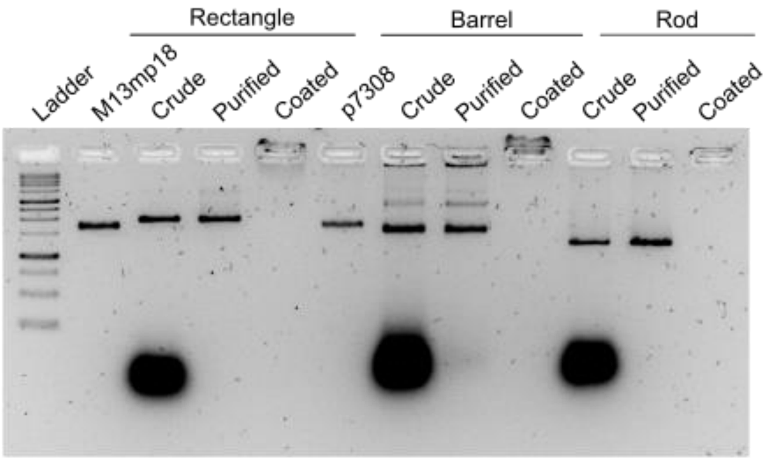


**S4.** Characterization of DN rectangle, barrel, and rods assessed through 2% agarose gel electrophoresis. The shift from the scaffold bands (M13mp18 and p7308) in the crude lane indicates DN folding. Purified lanes show the removal of excess staples by two rounds of PEG purification. K10-PEG coated structures are retained in the well. The gel was run for 2 hours on ice at 60V.

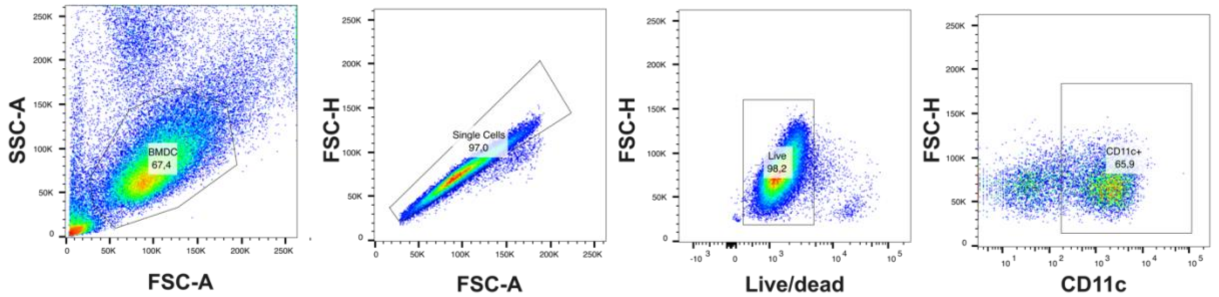


**S5.** Validation of BMDC differentiation using antibody staining. Bone marrow cells were cultured with GM-CSF for 7 days and then a control sample was stained with a viability dye and anti-CD11c. This gating strategy revealed that ~66% of the population were CD11c$^+$.

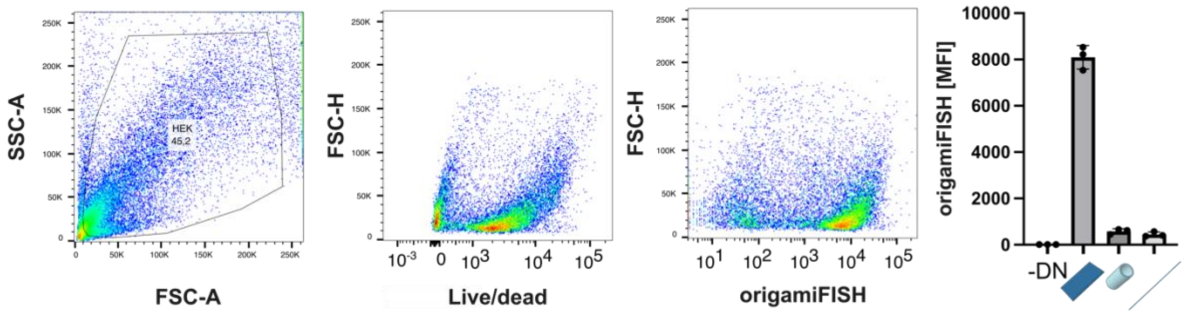


**S6.** The gating strategy of the HEK293 cells after uptake of DN shapes. (a) Forward scatter and side scatter profiles as well as the viability dye indicates a loss of a distinct live cell population. (b) Respective uptake of DN shapes from the gated cells using the strategy in (a).

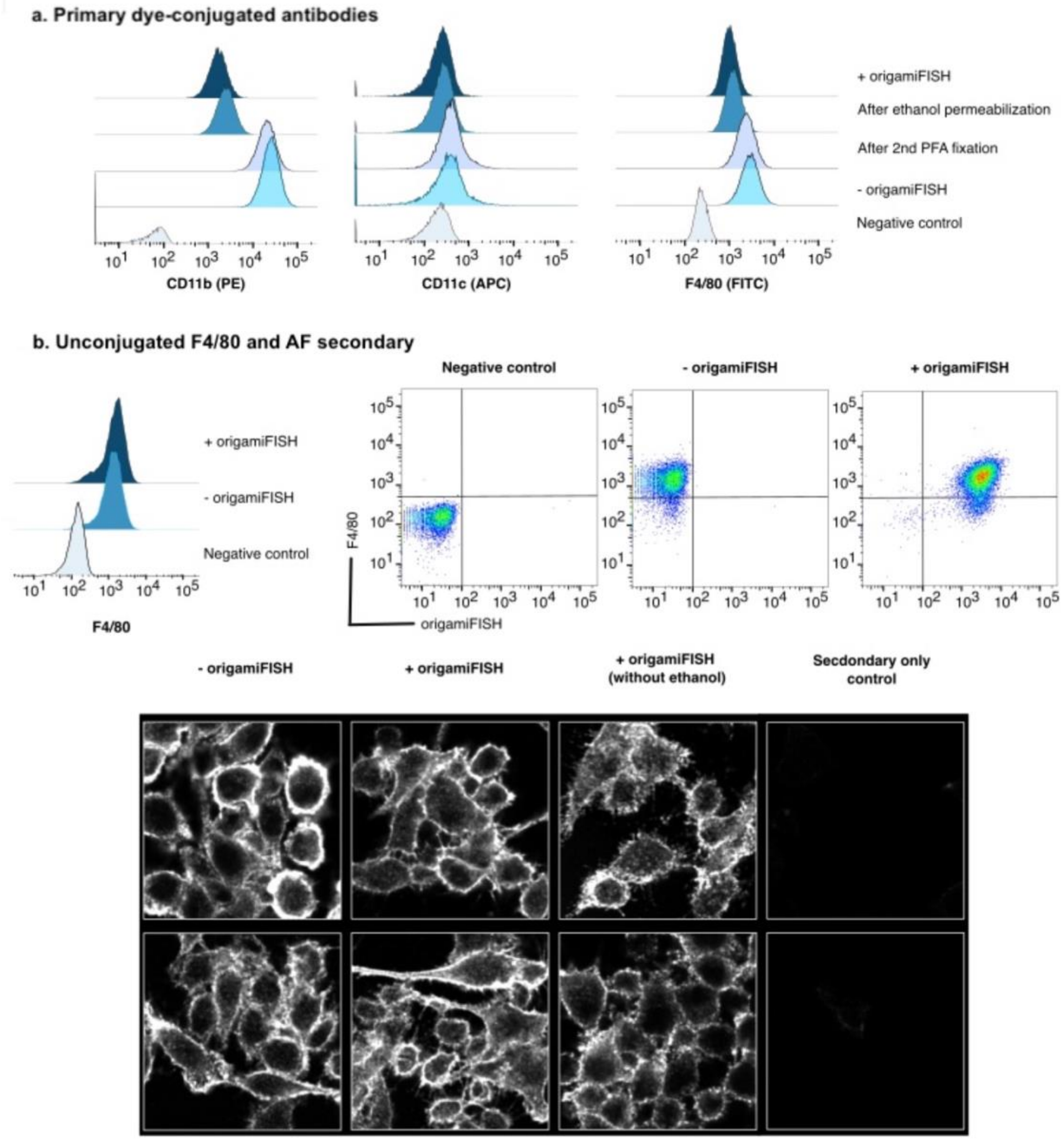


**S7.** Flow cytometry data of the signal loss using conjugated primary antibodies during origamiFISH-Flow protocol. (a) Raw 264.7 cells were stained for CD11b, CD11c, or F4/80 at varying steps of the origamiFISH protocol. (b) Flow cytometry data and confocal images of antibody signal retention though origamiFISH-Flow using unconjugated primary antibody F4/80 with Alexa 594 fluorophore-conjugated secondary antibody to label suggesting that the F4/80 epitope remains intact but the FITC dye in (a) is unable to persist.

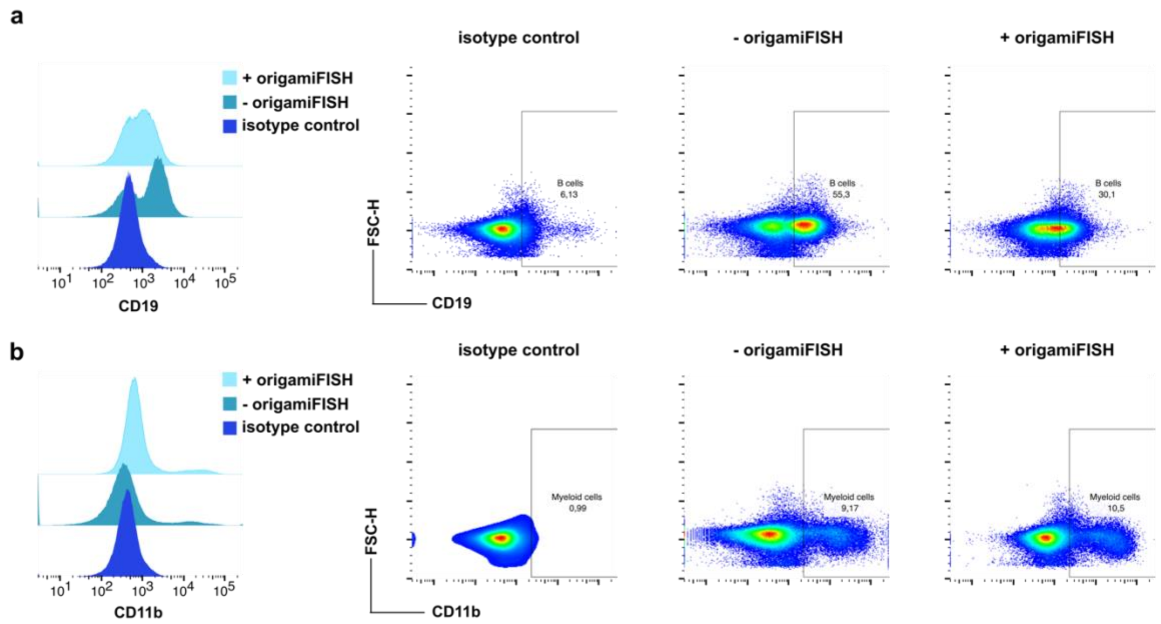


**S8.** Signal loss and retention of certain epitopes on splenocytes after the origamiFISH-Flow protocol. (a) origamiFISH-Flow caused $CD19^+$ populations to be undetectable but (b) retained the $CD11b^+$ population.

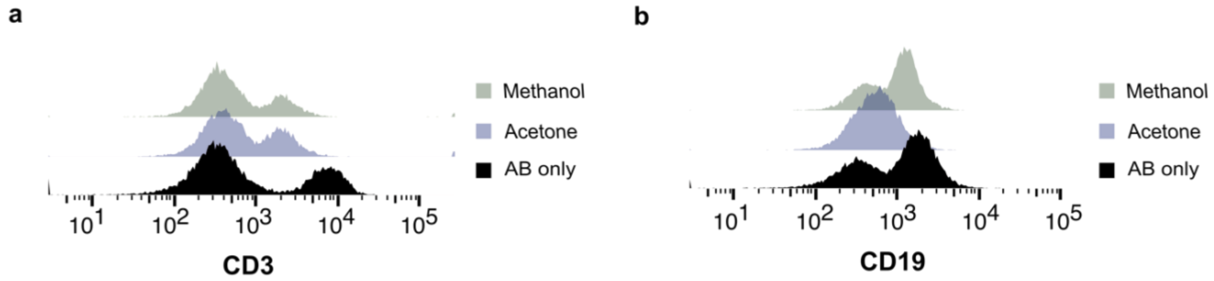


**S9.** Effects of replacing fixation and ethanol permeabilization steps with methanol or acetone as an alternative to ethanol for origamiFISH detection. Antibody (AB) only controls did not go through the origamiFISH protocol. Using acetone and methanol causes a loss of the $CD3^+$ population compared to the control. **CD3:** The AB only control stained 30% $CD3^+$ cells whereas, acetone detected 19% and methanol detected 16%. **CD19:** The AB only control and methanol stained 61% $CD19^+$ cells whereas, acetone caused a loss of the distinct positive population.

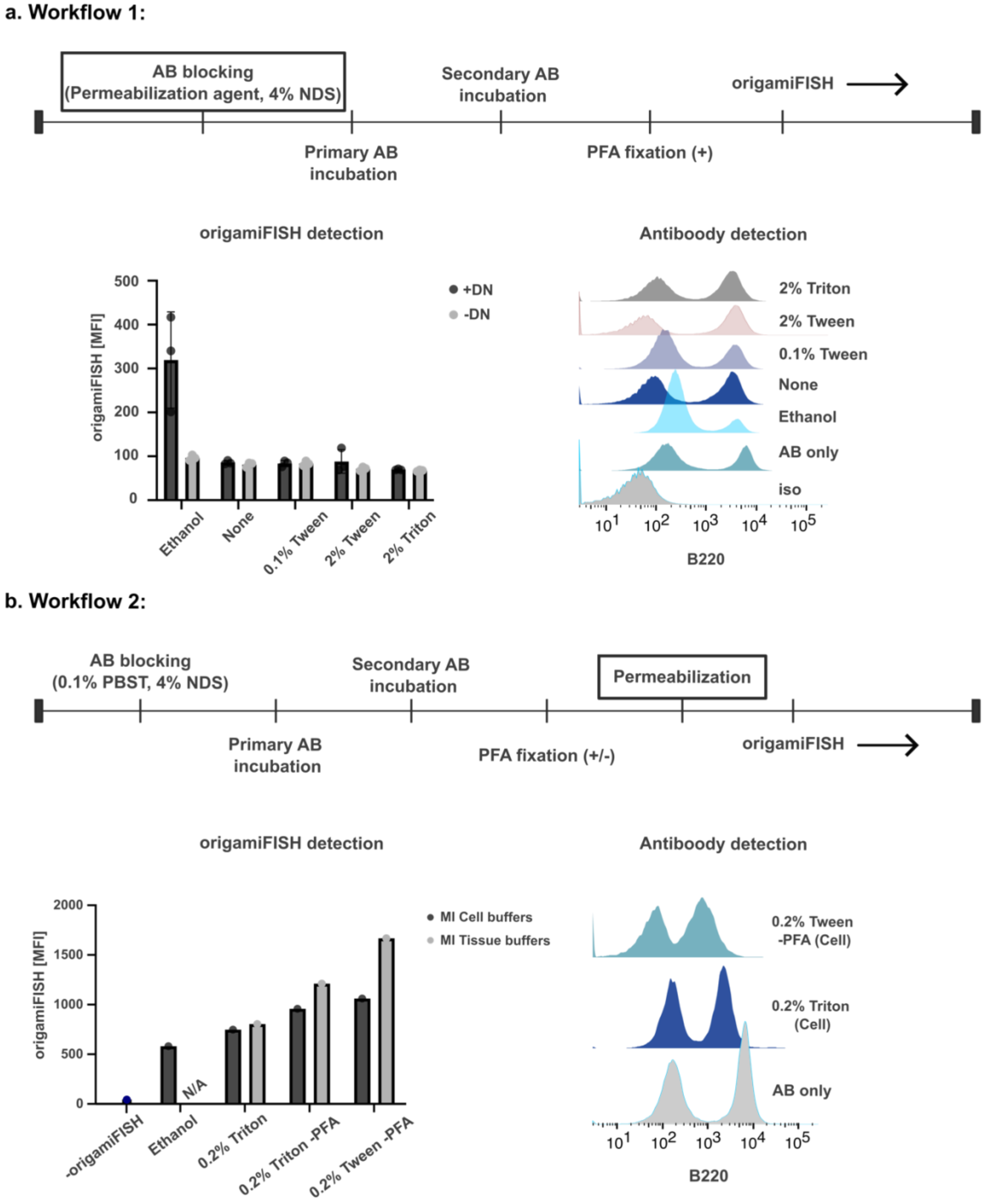


**S10.** Workflow optimization to combine antibody and origamiFISH-Flow staining protocols. (a) Workflow 1 consists of a permeabilization step within the antibody blocking. Tween 20 or Triton, at the specified concentration, was diluted in 4% normal donkey serum in 1X PBS while rocking for one hour. The AB only sample was blocked with 0.1% Tween 20 and did not proceed through origamiFISH. AB staining of B220 demonstrated a compatibility with Tween 20 and Triton but these reagents were inefficient at detecting origamiFISH signal. Ethanol treated cells were ineffective at retaining the entire B220 population but was the only condition that yielded effective origamiFISH detection. (b) In workflow 2, a 30-minute permeabilization with the mentioned reagents were done immediately prior to the origamiFISH protocol. Here, all conditions tested were able to detect origamiFISH signal. Moreover, using 0.2% Triton or 0.2% Tween 20 was able to retain B220 AB signal. MI, Molecular Instruments.

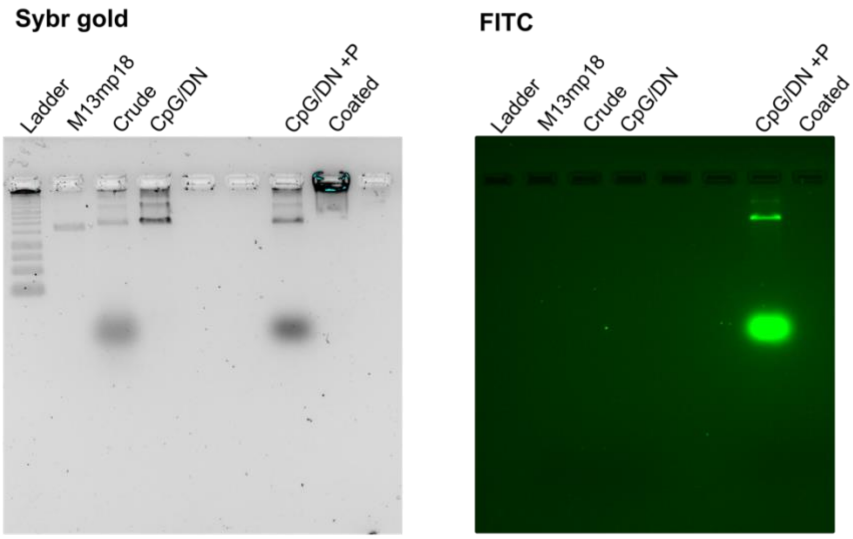


**S11.** 2% Agarose gel electrophoresis of CpG functionalized rectangle DN used for in vivo experiment. **SYBR Gold:** Band shift upwards from the crude lane to the M13mp18 scaffold indicates that the DNs folded. DNs were purified with PEG purification and functionalized with CpG (CpG/DN). **FITC:** FAM labeled anti-CpG probes were incubated for 1 hour with purified CpG/DN (CpG/DN +P). The second band in the CpG/DN + P lane is representative of excess probes. Purified and functionalized structures were coated with K10-PEG for 30 minutes at room temperature (Coated). The gel was run at 60V for 2 hours and then imaged on the FITC channel. Gel was then stained for 1 hour with SYBR gold and imaged on the universal channel. P, FAM labeled anti-CpG probe.
